# Characterizing flexibility and mobility in the natural mutations of the SARS-CoV-2 spikes

**DOI:** 10.1101/2021.09.14.460264

**Authors:** James Panayis, Navodya S. Römer, Dom Bellini, A. Katrine Wallis, Rudolf A. Römer

## Abstract

We use *in silico* modelling of the SARS-CoV-2 spike protein and its mutations, as deposited on the Protein Data Bank (PDB), to ascertain their dynamics, flexibility and rigidity. Identifying the precise nature of the dynamics for the spike proteins enables, in principle, the use of further *in silico* design methods to quickly screen for existing and novel drug molecules that might prohibit the natural protein dynamics. We employ a recent protein flexibility modeling approach, combining methods for deconstructing a protein structure into a network of rigid and flexible units with a method that explores the elastic modes of motion of this network, and a geometric modeling of flexible motion. Our results thus far indicate that the overall motion of wild-type and mutated spike protein structures remains largely the same.

## Introduction

In recent years, drug discovery supported by computer modelling has become a viable method to support experimental efforts in large-scale screening of prospective drug compounds [1]. This approach is part of the overarching theme of bioinformatics which aims to harness the power of modern computational techniques for “in silico” design of medicines [2]. The advantage of this lies not only in the increased screening speed, but also in its predictive power for novel drug development.

In early 2020, protein structures related to SARS-CoV-2 began to appear in the international Protein Data Bank (PDB). Such structures give the precise location of atoms in each protein, forming the basis for an understanding on how their biological mechanism might function. Chief among these structures were the protease (PDB code 6LU7 [3]), important in the life cycle of SARS-CoV-2, and the ectodomain of the spike glycoprotein (3 structures with PDB codes 6VSB [4], 6VXX and 6VYB [5]), a homo-trimer protein, protruding out of the viral capsid, which the virus uses in attaching itself to a human cell during infection. The spike protein is hence a main target for human antibodies and a key molecule for vaccine design and possible therapeutic treatments. The spike is a homo-trimer, protruding from the viral surface. The ectodomain of each monomer consists of an N-terminal subunit, S1, comprising two domains, S^A^ or N-terminal domain (NTD) and S^B^ or Receptor-Binding Domain (RBD), followed by an S2 subunit forming a stalk-like structure. Each monomer has a single membrane-spanning segment and a short C-terminal cytoplasmic tail [6].

In March 2021, Gobeil *et al*. [7] reported the first structures for the mutations of SARS-CoV-2 emerging widely in the United Kingdom (*α*-variant), South Africa (*β*-variant) and Brazil (*γ*-variant), as well as for the variant identified in the Danish mink population. All these variants either retain or improve binding to human cells [7].

In order to understand the biological function of a given structure such as the spike protein, it is important to know how such a molecule moves. Experimentally, this is very hard to ascertain and, even if achievable in some cases, may require months of careful experimentation. Computational modelling, on the other hand, can provide crucial information faster. However, in making such models, one has to make simplifying assumptions that can reduce the accuracy of the results that are obtained. In our work [8], we used an approach in which a protein structure is divided into rigid and flexible parts. This then allows us to quickly model the motion of the protein by keeping the rigid parts stable whilst allowing movement for the flexible regions.

In the following, we shall briefly explain the details of the computational modelling used and review the results obtained for the original “wild-type” spike proteins, which are only a small subset of the nearly 300 SARS-CoV-2 related protein structures studied in [8]. In addition, the same study was extended to the available PDB structures of variants, namely *α*, *β* and *γ*, which have emerged through various mutations [7].

## Methods

### Protein selection

The wild-type structures of the spike protein were deposited into the PDB already in March 2020 with code 6VSB [4] followed shortly afterwards by 6VXX and 6VYB [5]. With RMS resolution of 3.5 Å, 6VSB has a slightly lower resolution than 6VXX at 2.8 Å and 6VYB at 3.2 Å and we shall mainly refer to 6VXX and 6VYB when mentioning the “wild-type” structures. These latter two structures are distinguished by 6VXX having all 3 S^B^ domains in a “closed” or “down” configuration, while 6VYB shows one of these in an “open” or “up” configuration [4]. S1 is involved in recognition of the human receptor angiotensin-converting enzyme 2 (ACE2) [6], eventually leading to viral entry [9].

PDB available structures of mutated spike proteins are the *α*-variant with 4 structures (codes 7LWS to 7LWV), the *β*-variant with 7 structures (codes 7LYK to 7LYQ) and the *γ*-variant (code 7LWW). The variants identified in the Danish mink population (9 PDB codes 7LWI to 7LWQ), giving broadly similar results, are not shown here due to space constraints.

### Coarse-grained protein flexibility dynamics

We start our rigidity, flexibility and mobility modelling with the given atomic coordinate files from the PDB in *.pdb* format. The details have been described previously in Refs. [10,11] and the particulars for the spike proteins in Ref. [8]. Briefly, the PDB structures are cleaned and hydrogenated, their rigid and flexible parts identified [12] then the lowest vibrational modes computed [13,14] and geometric simulation along the simplest non-trivial modes is performed [15]. This explores the flexible motion available to a protein within a given pattern of rigidity and flexibility. Typically, several thousand new conformers for each mode are generated unless or until prohibited by steric clashes and/or bonding constraints. We emphasize that the resulting trajectories do not represent actual stochastic motion in a thermal bath as a function of time, but rather the *possibility of motion* along the most relevant elastic modes. The usefulness of this approach has previously been shown in a variety of systems [11, 16–19]. Methods similar in spirit, exploring flexible motion using geometric simulations biased along “easy” directions, have also been implemented by other groups [20,21].

## Results

### Rigidity and Flexibility

The rigidity pattern of the homotrimer in the wild-type structures 6VXX and 6VYB is dominated by a large rigid cluster encompassing most of the trimer structure [8], except for the region in one of the monomers of 6VYB corresponding to the RBD in the “up” or “open” configuration (Fig. 1). Hence the rigidity clearly distinguishes open from closed structures [23]. This study has found that this pattern repeats itself for the mutated variant structures shown in Table 1.

**Figure 1.**
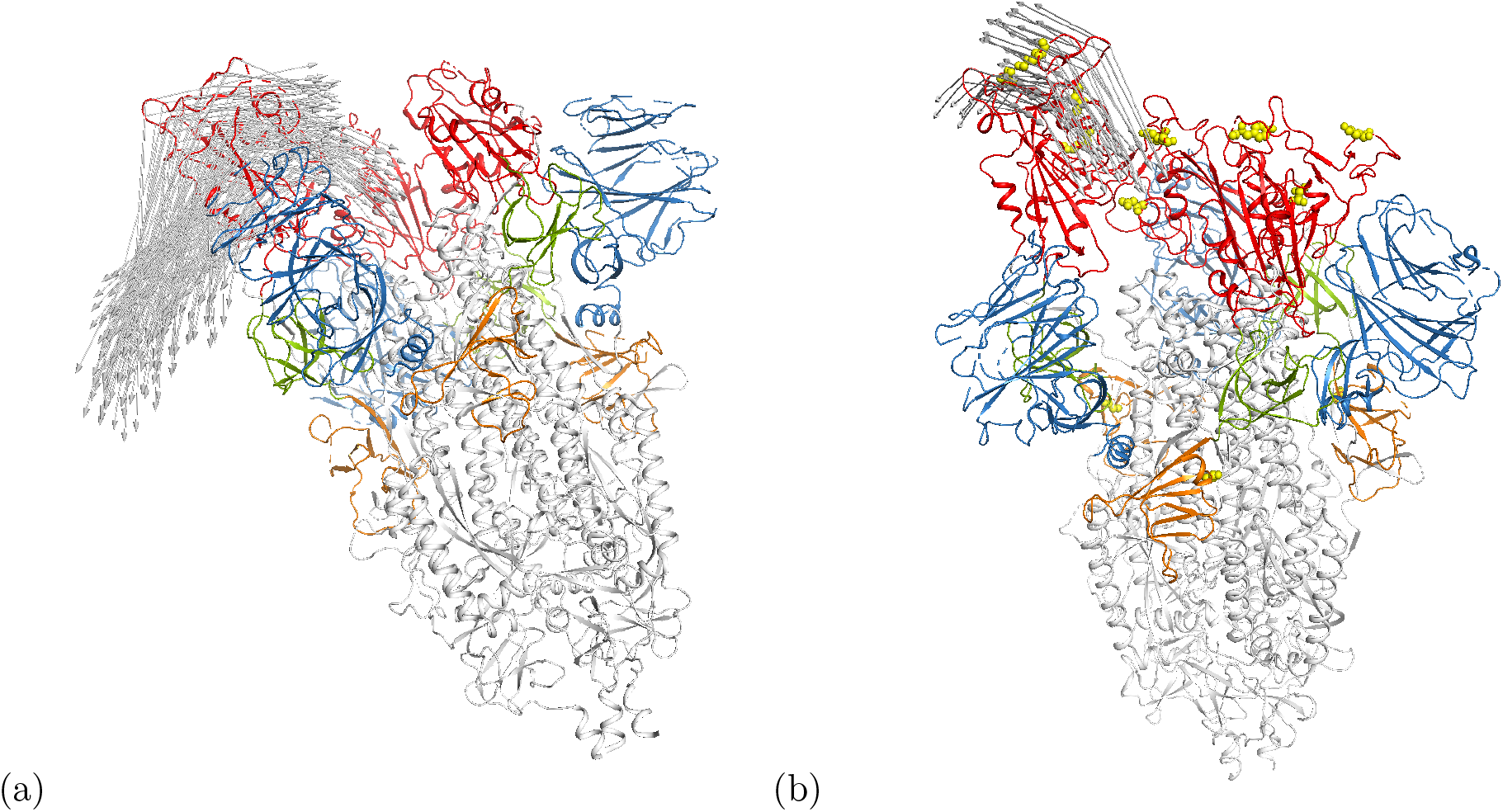
Cartoon representation of motion along modes *m*_7_ at *E*_cut_ = 1 kcal/mol in the (a) *open* wild-type spike ecto domain (6VYB) [5] and (b) the *γ*-variant 7LWW. Arrows indicate motion distances larger than 17.5Å. The RBDs are shown in red while the positions of mutations for 7LWW are given by the yellow spheres. Some other domains are also highlighted in color, i.e. the NTD is blue. An interactive animation of the motion is available for 6VYB [24] and 7LWW [25].

**Table 1.** List of SARS-Cov-2 spike structures investigated in this study. In column 3, a rigidity dilution classification into solid “brick”, 1st and 2nd order behaviour, and rigidity domains is given [8]. In column 5, the use of descriptor “flexibility” indicates largely elastic behaviour while “motion” denotes larger scale movements.

| PDB | Ref. | rigidity class. | mutation | behaviour at $\vert E_{\text{cut}}\vert = 1$ | u/d |
| --- | --- | --- | --- | --- | --- |
| 6VXX | [5] | domain | wild-type | $m_7$: flexibility in NTD-A/NTD-C, $m_8$: flexibility in NTD-B | 3 down |
| 6VYB | [5] | domain | wild-type | $m_7$: motion in RBD-B, $m_8$: motion in RBD-B, NTD-C | 1 up |
| 7LWS | [7] | 1st | $\alpha$ (UK) | $m_7$: flexibility in RBD-A+C, NTD-B, $m_8$: flexibility in NTD-A+C, RBD-A+C | 3 down |
| 7LWT | [7] | 1st | $\alpha$ (UK) | $m_7$: motion in RBD-B, $m_8$: flexibility in RBD-B, NTD-C | 1 up |
| 7LWU | [7] | 1st, domain | $\alpha$ (UK) | $m_7$: motion in RBD-B, $m_8$: motion in RBD-B | 1 up |
| 7LWV | [7] | 1st, domain | $\alpha$ (UK) | $m_7$: motion in RBD-B, $m_8$: motion in RBD-B, flexibility in NTD-C | 1 up |
| 7LYK | [7] | 1st, domain | $\beta$ (SA) | $m_7$: motion in RBD-B, $m_8$: motion in RBD-B+A | 2 up |
| 7LYL | [7] | 1st | $\beta$ (SA) | $m_7$: flexibility in NTD-A, RBD-B+C, $m_8$: flexibility in RBD-A, NTD-B+C | 3 down |
| 7LYM | [7] | 1st | $\beta$ (SA) | $m_7$: flexibility in NTD-A, RBD-B+C, $m_8$: flexibility in RBD-A, NTD-B+C | 3 down |
| 7LYN | [7] | 1st | $\beta$ (SA) | $m_7$: motion in RBD-B, $m_8$: motion in RBD-B, NTD-C | 1 up |
| 7LOY | [7] | 1st, domain | $\beta$ (SA) | $m_7$: motion in RBD-B, $m_8$: flexibility in NTD-C | 1 up |
| 7LYP | [7] | 1st | $\beta$ (SA) | $m_7$: motion in RBD-B, $m_8$: motion in NTD-C, flexibility in RBD-B | 1 up |
| 7LYQ | [7] | 2nd | $\beta$ (SA) | $m_7$: motion in RBD-B, $m_8$: motion in RBD-B, NTD-C | 1 up |
| 7LWW | [7] | 1st | $\gamma$ (BR) | $m_7$: motion in RBD-B, $m_8$: motion in NTD-C, flexibility in RBD-B | 1 up |

### Protein Mobility

We have performed a Froda mobility simulation for each protein at several selected values of *E*_cut_. In a previous publication, we discussed the criteria for a robust selection of *E*_cut_ [15]. For each protein structure as given in Table 1, we identified a bespoke set of *E*_cut_ values from the rigidity plots corresponding to values of important loss of rigidity. In Fig. 1, we choose *E*_cut_ = –1 kcal/mol and show results for the simplest non-trivial mode, *m*_7_, with 6VYB and 7LWW. Clearly, 6VYB shows good motion in the RBD domain for monomer B. This is similar to the mutated structure 7LWW. In addition, we also find for 7LWW [25] that more motion is visible in the N-terminal domain (NTD) on monomer C than in the NTDs of 6VYB [24].

In Fig. 2 we quantify the motion by showing, for modes *m*_7_ and *m*_8_ of 6VYB and 7LWW, the root-mean-squared deviation (RMSD) compared to the initial (crystal) structure along the motion trajectories. For *m*_7_, the motion of the RBD domain on monomer B can be clearly seen to emerge around conformer 50, corresponding to around 1000 steps in the motion simulation. Somewhat later, one sees the RMSD increase for NTD C on 7LWW. Mode ms has an even richer pattern of behaviour with nearly as much RMSD in NTD C as in RBD B in both structures.

**Figure 2.**
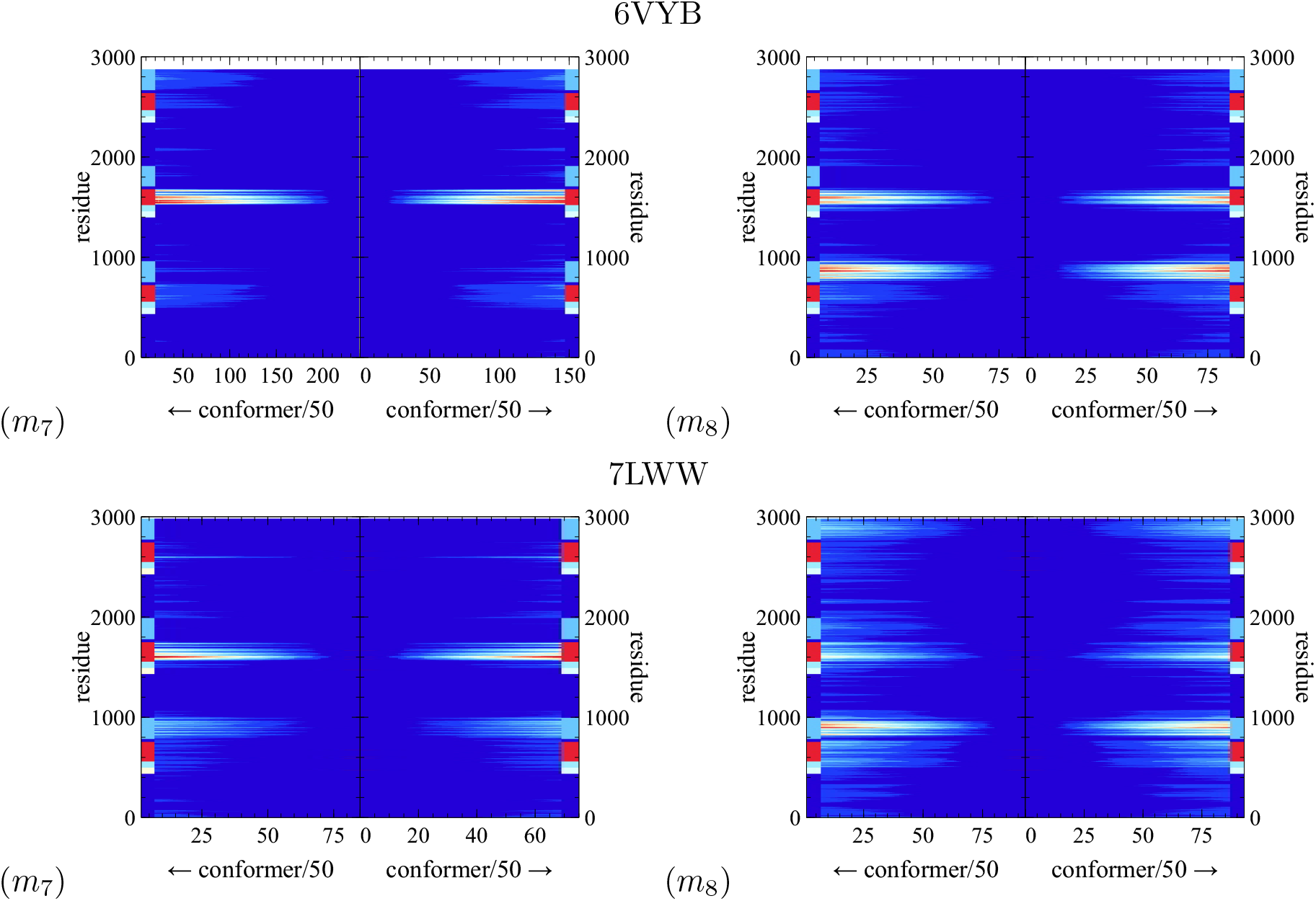
RMSD variation along modes *m*_7_ (top) and *m*_8_ (bottom) at *E*_cut_ = 1 kcal/mol in the (a) *open* spike ecto domain (6VYB) [5] and (b) the *γ*-variant 7LWW. The colored blocks at the left and right in each panel indicate domains with the RBD given in red while the NTD is cyan.

Similar results have been obtained for the other mutations listed in Table 1. For modes 7 and 8 at *E*_cut_ = 1 kcal/mol, the average RMSD across both modes and both directions (and the average of the largest observed RMSDs of each mode) for wild-types (6VXX, 6VYB), *α* (7LWS – 7LWV), *β* (7LYK – 7LYQ), and γ (7LWW) variants are respectively: 2.53Å (18.02Å), 2.95Å (22.99Å), 2.1Å (15.44Å) and 2.28Å (17.67Å). We find that, naturally, closed spike structures exhibit a smaller RMSD than the open structures. In addition, open structures seem to have roughly the same change in RMSD during the movement. Overall, this suggests that with respect to our modelling, there is no statistically relevant distinction between wild-type and mutated structures or between the mutated structures themselves.

## Conclusions

The mutations of the SARS-CoV-2 virus that have emerged thus far in the human population are largely concerned with a few isolated mutations of residues on the spike protein. It seems entirely plausible that such isolated mutations should not dramatically change the overall flexibility of a protein with nearly 3000 residues. This is what we observe. Since the spikes are basically the prime functional target for all existing vaccines, our results provide reassurance that, at least from a geometric simulation approach, the geometry of the vaccines needs no major readjustment to continue to work also for the mutations of the spikes. However, since the mutations nevertheless are known to have somewhat different responses to vaccines, our result might also suggest that factors beyond simple geometric modelling are important – with Coulombic surface charges on proteins clearly being among the main suspects. On the other hand, the structures as used here are taken as deposited in the PDB. Perhaps an equilibration and thermalization process might lead to structural refinements that could also cause conformation changes in the motion trajectories as computed here. Investigations along these lines are proceeding.

## Acknowledgements

We thank Young-Jun Park for pointing us to Ref. [7]. JP and RAR thankfully acknowledge funding via Warwick’s Undergraduate Research Support Scheme (URSS). We thank Warwick’s Scientific Computing Research Technology Platform for computing time and support. UK research data statement: Data accompanying this publication are available for download at [22].

